# Lipid peroxidation increases membrane tension, Piezo1 gating and cation permeability to execute ferroptosis

**DOI:** 10.1101/2022.10.31.514557

**Authors:** Yusuke Hirata, Ruiqi Cai, Allen Volchuk, Benjamin E. Steinberg, Atsushi Matsuzawa, Sergio Grinstein, Spencer A. Freeman

**Author notes:** Corresponding authors: Spencer A. Freeman and Sergio Grinstein.

## Abstract

The ongoing metabolic and microbicidal pathways that support and protect cellular life generate potentially damaging reactive oxygen species (ROS). To counteract damage, cells express peroxidases, antioxidant enzymes that catalyze the reduction of oxidized biomolecules. Glutathione peroxidase 4 (GPX4) is the major hydroperoxidase specifically responsible for reducing lipid peroxides; this homeostatic mechanism is essential and its inhibition causes a unique type of lytic cell death, ferroptosis. The mechanism(s) that lead to cell lysis in ferroptosis, however, are unclear. We report that the lipid peroxides formed during ferroptosis accumulate preferentially at the plasma membrane. Oxidation of surface membrane lipids increased tension on the plasma membrane and led to the activation of Piezo1 and TRP channels. Oxidized membranes thus became permeable to cations, ultimately leading to gain of cellular Na^+^ and Ca^2+^ concomitant with loss of K^+^. These effects were reduced by deletion of Piezo1 and completely inhibited by blocking cation channel conductance with ruthenium red or 2-aminoethoxydiphenyl borate (2-APB). We also found that the oxidation of lipids depressed the activity of the Na^+^/K^+^-ATPase, exacerbating the dissipation of monovalent cation gradients. Preventing the changes in cation content attenuated ferroptosis. Together, our study establishes that increased membrane permeability to cations is a critical step in the execution of ferroptosis and identifies Piezo1, TRP channels and the Na^+^/K^+^-ATPase as targets/effectors of this type of cell death.

## Introduction

Programmed cell death or apoptosis, was first described a half century ago ^1^. Since then, multiple modes of non-apoptotic cell death have been identified and characterized including necroptosis, pyroptosis, and most recently, ferroptosis. Each of these pathways is governed by a combination of unique and overlapping molecular mechanisms, though the precise pathways leading to ferroptosis are the least understood ^2^.

Ferroptosis is a caspase-independent form of cell death initiated by the production of Fe^2+^-dependent accumulation of lipid peroxides ^3^. Lipid peroxides are in fact continuously formed by organellar-derived reactive oxygen species (ROS) that combine with reactive hydrogens in the polyunsaturated acyl chains of membrane lipids ^4^. Excessive ROS production, for example from mitochondria, can contribute to the membrane damage that underlies a range of pathologies ^5^. Thus, lipid peroxidation must be highly reversible and tightly controlled ^6^. The major enzyme responsible for such control is glutathione peroxidase 4 (GPX4), which reduces toxic phospholipid hydroperoxides to non-toxic phospholipid alcohols at the expense of glutathione (GSH). GPX4 thereby prevents the accumulation of oxidized lipids and ultimately, ferroptotic cell death ^7^. Accordingly, ferroptosis can be induced directly by the inhibition of GPX4 or indirectly by depleting GSH^3, 7^.

ROS production varies between cell types depending on the metabolic and microbicidal pathways they employ. The ROS produced by mitochondria are related to the number of potential electron donors in the transport chain and oxygen concentration: when mitochondria are not making ATP, have a high H^+^-motive force and a diminished pool of coenzyme Q, ROS production increases ^8^. Glycolytic cancers or those in hypoxic conditions are therefore highly susceptible to ferroptosis ^9^. This susceptibility has prompted high throughput screening for compounds that may serve as selective GPX4 inhibitors that may preferentially target transformed cells. These efforts resulted in the identification of RAS-selective lethal 3 (RSL3) ^7^.

In addition to being considered as a means for tumor suppression, ferroptosis is also associated with degenerative diseases of various organs (e.g. the brain, heart, liver, and kidney), collectively referred to as ferroptosis-related diseases ^6^. In these instances, the use of ferrostatin-1 (Fer-1) ^3^ –a synthetic antioxidant that scavenges alkoxyl radicals– to supplement the activity of GPX4 has been proposed. Clearly, ferroptosis has emerged as a tunable and targetable pathway involved in the pathogenesis of various important diseases.

Despite the impact of ferroptosis on health, precious little has been described regarding the connections between lipid peroxidation and cell death, or even the precise site(s) of lipid peroxide accumulation in the cell. Other forms of cell death are associated with dysregulation of the ionic composition of the cytosol and the attendant cell volume changes, intimating some possibilities. Apoptosis, for example, is characterized by cell shrinkage mainly due to loss of cytosolic K^+^ and osmotically obliged water, which consequently potentiate apoptotic cell death ^10^. In contrast, cells swell when undergoing pyroptosis, despite the loss of K^+^, leading to cell rupture ^10^. In spite of these well-characterized phenomena, there are no previous reports examining monovalent cation fluxes during ferroptosis and the possible occurrence of cell volume change has not been assessed directly, remaining the subject of controversy; it therefore remains unknown whether any such changes are required for death to occur.

In the present study, we sought to understand the role of ionic homeostasis in ferroptotic cell death. We report that the lipid peroxides that form upon the inhibition of GPX4 or GSH synthesis accumulate predominantly in the plasma membrane, increasing its tension and permeability to monovalent cations. Drastic changes in monovalent cation composition occur before the cells begin to rupture: a large Na^+^ gain together with a loss of K^+^, which are associated with an increase in the net cell volume. The cation content changes are mediated by the activation of mechanosensitive non-selective cation channels, piezo-type mechanosensitive ion channel component 1 (Piezo1) and the transient receptor potential (TRP) channel family. The effect is further exacerbated by the inactivation of the sodium-potassium (Na^+^/K^+^)-ATPase which is responsible for maintaining the concentration gradients of these cations across the plasma membrane in healthy cells. We could inhibit ferroptosis by preventing the changes in ionic content, delaying ferroptotic cell death without affecting the amount of oxidized lipid in the membrane, thus revealing an important parameter controlling ferroptosis.

## Results

### The lipid peroxides formed during ferroptosis preferentially accumulate in the plasma membrane preceding cell rupture

The inhibition of GPX4 by RSL3 leads to ferroptosis across various cells and tissues. We initially examined the dose-response and time course of cell rupture induced by RSL3 to establish suitable experimental conditions to interrogate the events that precede cell lysis. Three cell lines of varying origin were used for this purpose: HT1080, RAW 264.7 (RAW), and HeLa. We found that after 24 h of treatment with RSL3 the maximal release of LDH –a good indicator of cell rupture– was achieved with 0.5 μM in HT1080 cells, 1.25 μM in RAW cells, and 5 μM in HeLa cells (Fig 1a).

**Figure 1.**
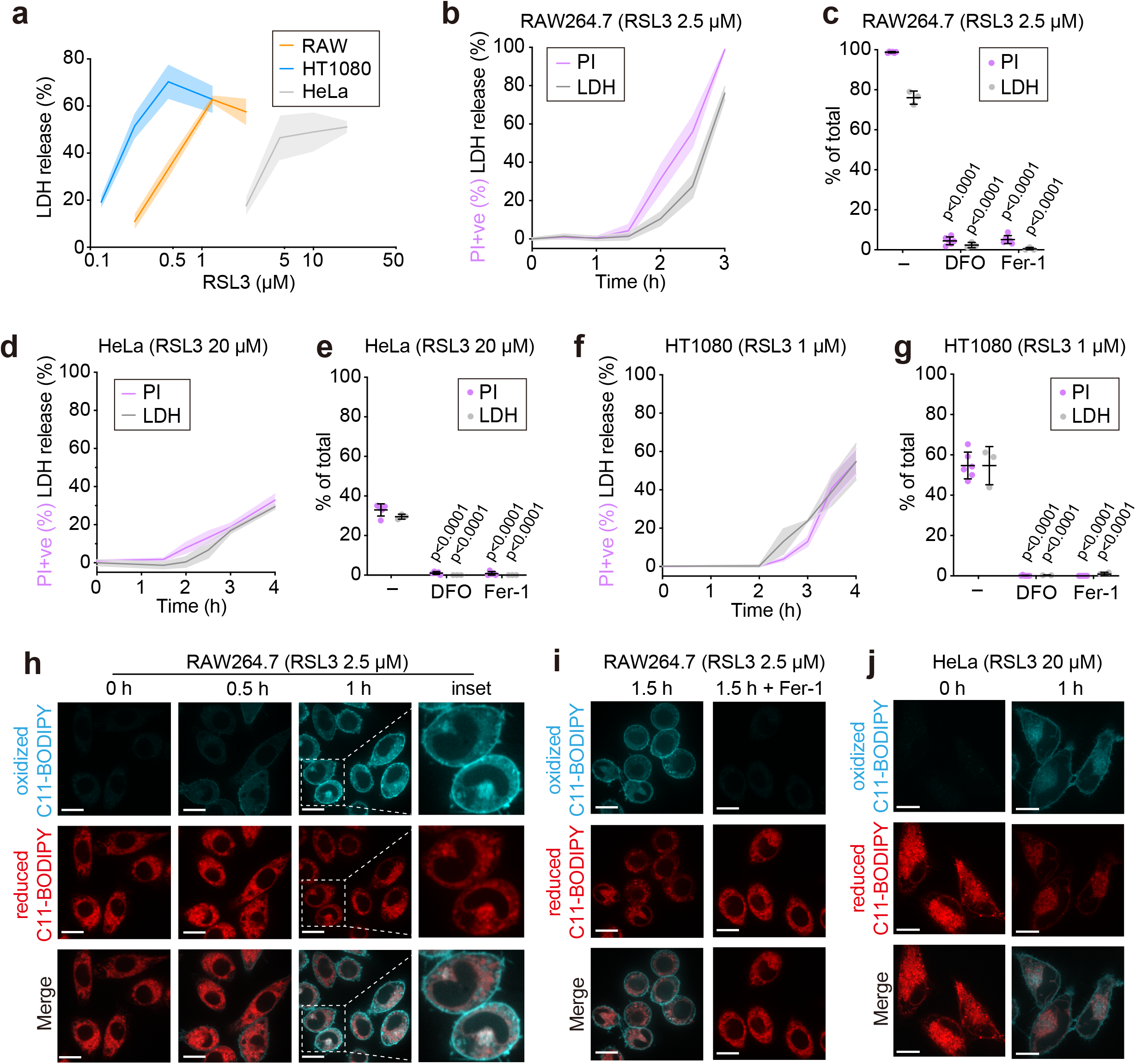
Lipid peroxides accumulate preferentially in the plasma membrane at the initiation of ferroptosis. a-g) LDH release and PI positivity (% of total) for RAW264.7, HeLa, and HT1080 cells. In *a*, cells were treated with indicated concentrations of RSL3 for 24 h. Cells were treated for indicated time periods in *b, c*, and *f*. In *c, e*, and *g*, 100 μM DFO or 2.5 μM Fer-1 were given at the time of RSL3 treatment for 3 h (RAW264.7) or 4 h (HeLa or HT1080 cells). In all cases, data are shown as mean ± SD, n=3. h-j) C11-BODIPY 581/591 imaging in RAW264.7 (h and i) and HeLa cells (j) treated with RSL3 +/-2.5 μM Fer-1 for indicated periods. Scale bars, 10 μm.

We next monitored the time-course of cell death. When treated with RSL3 at 2.5 μM, RAW cells were ∼30% positive for propidium iodide (PI) after 2 h and 100% positive after 3 h (Fig 1b,c); with 20 μM RSL3 HeLa were 30% of PI-positive after 4 h (Fig 1d,e); with 1 μM of RSL3 HT1080 cells were 50% PI-positive after 4 h (Fig. 1f,g). LDH release tracked closely with PI positivity in all cases (Fig 1b-g). Importantly, ferroptosis was completely reversed when RSL3 was delivered together with either the iron chelator deferoxamine (DFO) or the scavenging antioxidant ferrostatin-1 (Fer-1; Fig. 1c,e,g). These experiments established the conditions used throughout most subsequent experiments in this study.

We next used the ratiometric lipid peroxidation sensor, C11-BODIPY 581/591, to determine the site(s) of lipid peroxide accumulation in live cells^11^. Oxidation of the polyunsaturated butadienyl portion of the C11-BODIPY 581/591, which partitions into lipid bilayers throughout the cell, causes a shift in its fluorescence emission peak from ∼590 nm (red) to ∼510 nm (pseudo-colored cyan in the images shown). As illustrated in Fig. 1h and 1i, oxidized C11-BODIPY became detectable as early as 0.5 h after the addition of RSL3. After 1 h, oxidized lipids were most apparent on the plasma membrane in RAW, HeLa, and HT1080 cells (Fig. 1h,j; Supplementary Fig. 1a,b), being less abundant in endomembranes. The specificity of the C11-BODIPY 581/591 probe was confirmed by treating the cells with Fer-1, which entirely prevented the appearance of the oxidized C11-BODIPY 510 nm signal (Fig 1i and Supplementary Fig. 1a,b). Thus, lipid peroxides were found to accumulate preferentially in the plasma membrane of cells exposed to RSL3 before their rupture (see 1-2 h time points in Fig 1b,d,f). Consistent with this finding, lipid peroxides were also observed in the plasma membrane upon treating the cells with ML210, another covalent inhibitor of GPX4^12^, or with L-buthionine sulfoximine (BSO), an inhibitor of GSH synthesis (Supplementary Fig. 1c). Taken together, the accumulation of lipid peroxides in the plasma membrane that precedes lytic cell death appears to be a common phenomenon in ferroptosis, independently of the means by which it is induced.

### NINJ1 is not required for lytic cell death associated with ferroptosis

A recent study revealed that cell rupture associated with most forms of cell death, including apoptosis and pyroptosis, is facilitated by the multimerization of a transmembrane protein, nerve injury-induced protein 1 (NINJ1)^13^. The role of NINJ1 in ferroptosis has not been tested. To this end, we edited RAW cells expressing the apoptosis-associated speck-like protein containing a CARD (RAW-ASC) with CRISPR/Cas9 to delete NINJ1 (Fig. 2a). We first confirmed that the NINJ1^-/-^ cells were refractory to cell rupture during pyroptosis (induced by LPS priming followed by nigericin treatment) and apoptosis (induced by either H_2_O_2_ or staurosporine) (Fig. 2b), consistent with the previous report^13^. However, the NINJ1^-/-^ cells remained susceptible to ferroptosis, undergoing rupture to the same extent as non-edited wildtype (WT) cells. This result suggests that NINJ1 does not play a role in lytic cell death during ferroptosis.

**Figure 2.**
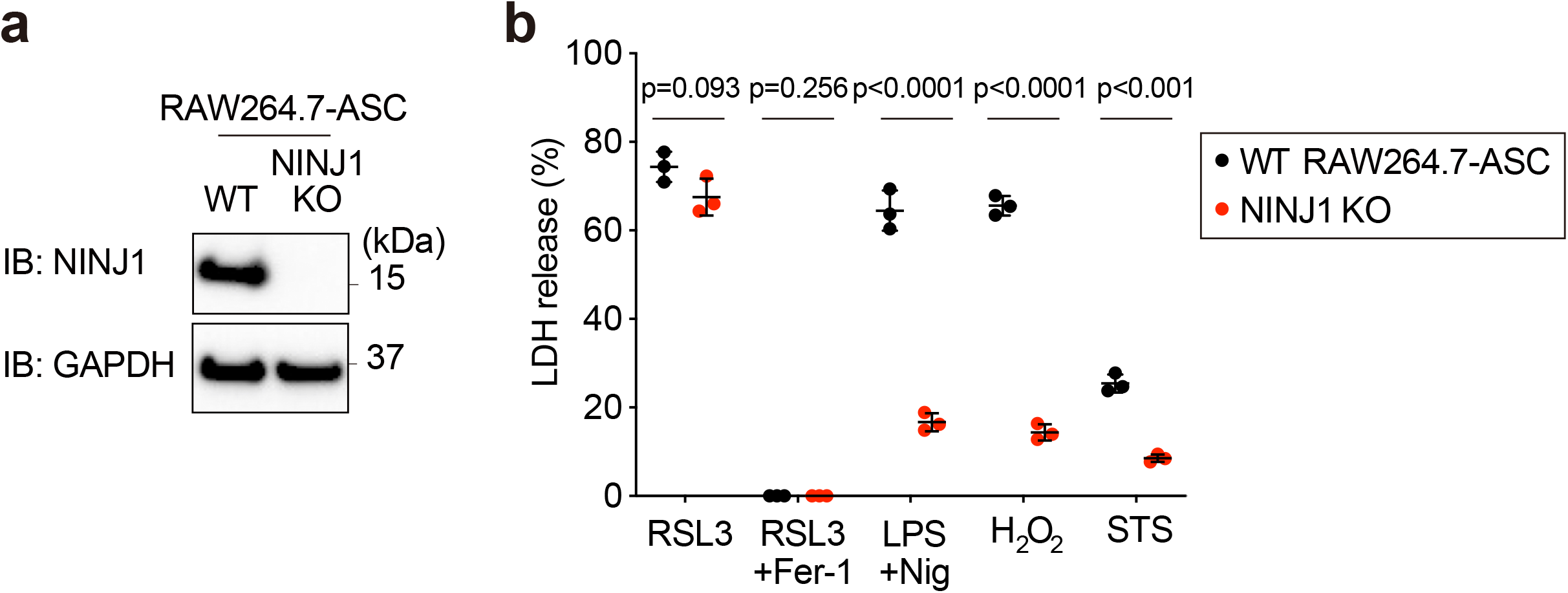
NINJ1 is not required for cell rupture during ferroptosis. a) Immunoblot for NINJ1 in WT and NINJ1 KO RAW-ASC cells. b) LDH release after 2.5 μM RSL3 treatment +/-2.5 μM Fer-1, LPS plus Nigericin (Nig) (cells were pretreated with 0.5 μg/mL LPS for 3 h, followed by 10 μM Nig treatment for 2 h), 400 μM H_2_O_2_ or 600 nM staurosporine (STS) for 5 h. Data are shown as the mean ± SD, n=3.

### Cell volume and plasma membrane tension increase before cell rupture

Ferroptosis has been associated with cell rounding and diminution of overall cell length, which has led to the assumption that the process involves volume loss^7, 14^. As the cells become round, however, they likely retract as they detach from the basolateral adhesive surface; as such, the shape change is not necessarily indicative of reduced volume. A more formal and rigorous quantitative determination of cell volume changes in cells undergoing ferroptosis was therefore warranted. In line with previous observations, RAW cells treated with the GPX4 inhibitor RSL3 became rounded within 0.5 h, as judged both by bright field imaging (Fig. 3a) and by visualization of the fluorescence of C11-BODIPY 581/591 (Fig. 1h,j). To determine if cell rounding was associated with changes in cell volume we used electronic cell sizing, an approach that accurately determines the volume of cells by the impedance to current as they flow through an aperture^15^. We found that, in contrast to previous assumptions, cell volume in fact increased in cells treated with RSL3 (Fig. 3b,c). The gain in cell volume ranged between 10-20% in RAW and HeLa cells and was not observed in cells treated with Fer-1 (Fig. 3b,c), suggesting that this effect was dependent on the lipid peroxidation. Mechanically lifting the cells could cause them to break or change their shape, potentially affecting their volume; we therefore validated the occurrence of volume changes in adherent cells by live-cell 3D imaging. To this end, cells were loaded with the fluorescent cytosolic dye CFSE and were subsequently imaged by spinning disc confocal microscopy. As shown in Fig. 3d, RSL3 treatment increased cell height and decreased undulations of the membrane. Quantification of the 3D images reconstructed from serial confocal slices using Volocity and Imaris software replicated an increase in cell volume upon RSL3 treatment (Fig. 3e), consistent with the results of electronic sizing (Fig. 3b).

**Figure 3.**
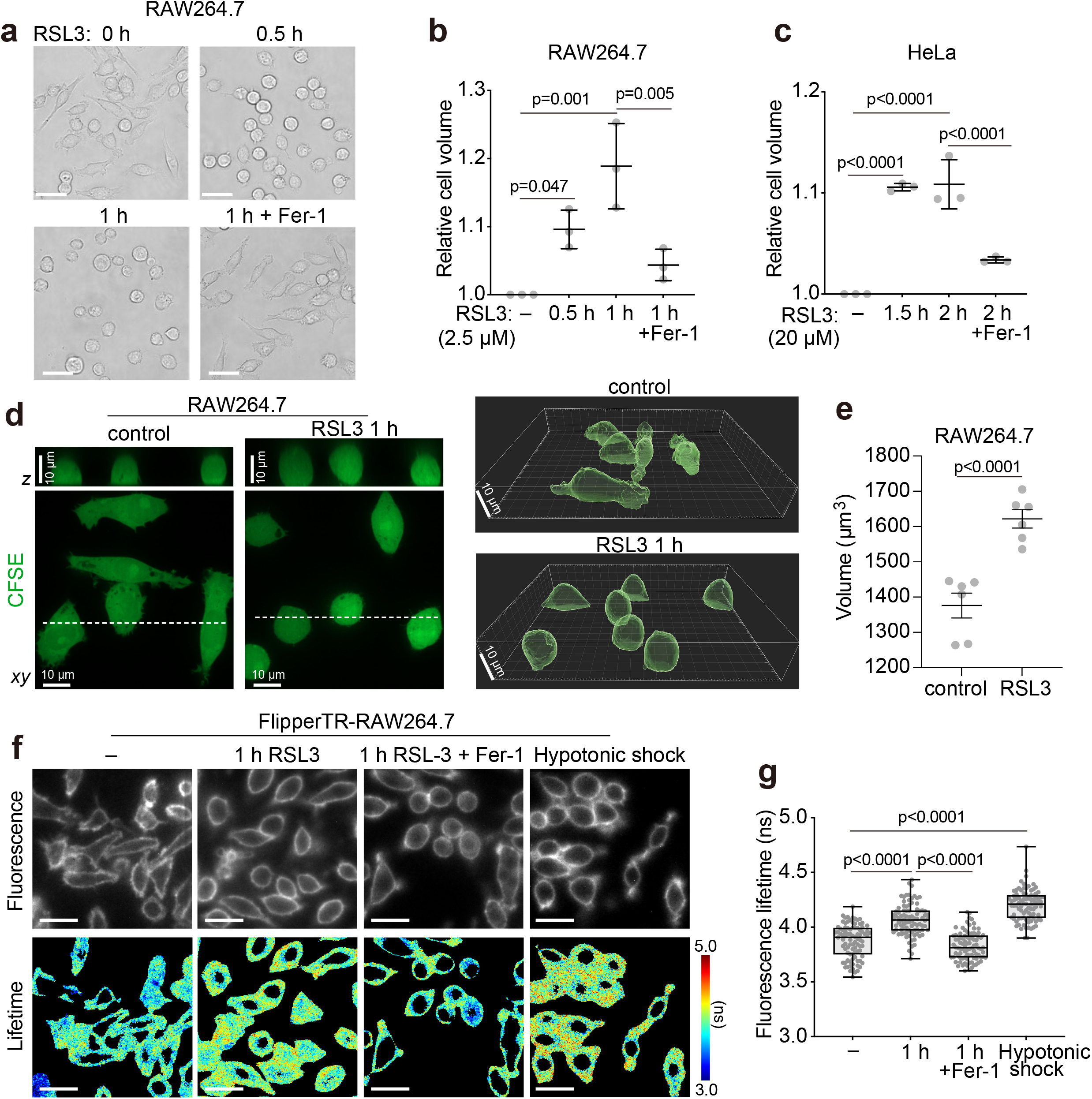
Lipid peroxidation leads to cell volume increases and higher plasma membrane tension. a) Bright field images of cells treated with 2.5 μM RSL3 +/-2.5 μM Fer-1 over time. Scale bar, 25 μm. b-c) Cell volume determinations for RAW264.7 (b) and HeLa cells (c) treated with RSL3 in the presence or absence of 2.5 μM Fer-1. The relative cell volume, mean ± SD, is shown. n=3. d-e) Cell volume determinations for RAW264.7 cells by live cell imaging. Images of control cells or cells treated 2.5 μM RSL3 for 60 min are shown in (d). The mean cell volume quantified per cell is indicated in *e*. n=6, >100 cells per condition. f-g) FLIM imaging of Flipper-TR in cells treated as in (a) or control cells in hypotonic (∼100 mOsm) solution for 2-3 min. Representative fluorescence images and heatmaps of fluorescence lifetime (color bar: lifetime in nanoseconds (ns)) are shown in (f). Scale bar, 25 μm. The mean lifetime of Flipper-TR quantified per cell is indicated in *g*. n=3, 90 cells per condition.

Shifts in cellular volume, uncompensated by a change in membrane surface area, are expected to alter plasma membrane tension^16^. Membrane tension can also increase due to expansion of its constituent lipids: modeling of phospholipid bilayers predicts that the bending of acyl chains that results from their oxidation will cause thinning and expansion of the membrane, elevating tension^17, 18^. The observation that ferroptosis was preceded by the accumulation of lipid peroxides in the plasma membrane, gross changes in cell shape and a concomitant increase in cell volume, motivated us to examine if these events were associated with changes in plasma membrane tension.

Flipper-TR^19^, which readily partitions into the plasma membrane and is sensitive to the changes in membrane tension, enables non-invasive measurements of membrane tension by fluorescence lifetime imaging microscopy (FLIM). Increases in the membrane tension prevent the rotation of the probe, increasing its fluorescence lifetime. We first analyzed the fluorescence lifetime in RAW cells before and after hypotonic shock (100 mOsm), which should cause a near-maximal increase in plasma membrane tension. Hypotonic shock increased the fluorescence lifetime by 0.33 ns (from 3.87 ns to 4.20 ns). RSL3 treatment of the cells for 1 h also increased the average lifetime (Fig. 3f,g). The increase in Flipper-TR fluorescence lifetime caused by RSL3 was completely eliminated in the presence of Fer-1 (Fig. 3f,g). Collectively, these results suggest that during ferroptosis, plasma membrane tension is elevated substantially.

### Piezo1 facilitates ferroptosis

Increases in membrane tension activate various mechanosensitive cation channels, including Piezo1^20^. Piezo1 is a non-selective cation channel (i.e. for Ca^2+^, Na^+^, K^+^, etc.) that opens in response to mechanical forces that deform the surface membrane, playing a critical role in various biological processes^21^. Interestingly, it has been reported that Ca^2+^ influx precedes cell rupture during ferroptosis^22^. In accordance with those observations, we indeed found that cytosolic Ca^2+^ –measured by expressing the genetically-encoded biosensor GCaMP6s– was markedly increased 90 min after addition of RSL3 (Fig. 4a), prior to cell rupture (Fig. 1b). To assess the role of mechanosensitive channels we treated cells with the broad cation channel inhibitor, ruthenium red (RR). Blocking these channels effectively inhibited LDH release instigated by various ferroptosis inducers including RSL3 (Fig. 4b), ML210 and BSO (Fig. 4c). Using DPPH (2,2-diphenyl-1-picrylhydrazyl) to assess antioxidant power, we determined that, unlike Fer-1, ruthenium red displays no antioxidant activity (Fig. 4d). Given that ruthenium red is cell-impermeant, these results are instead consistent with the notion that it protects cells from ferroptosis by blocking cation channel activity.

**Figure 4.**
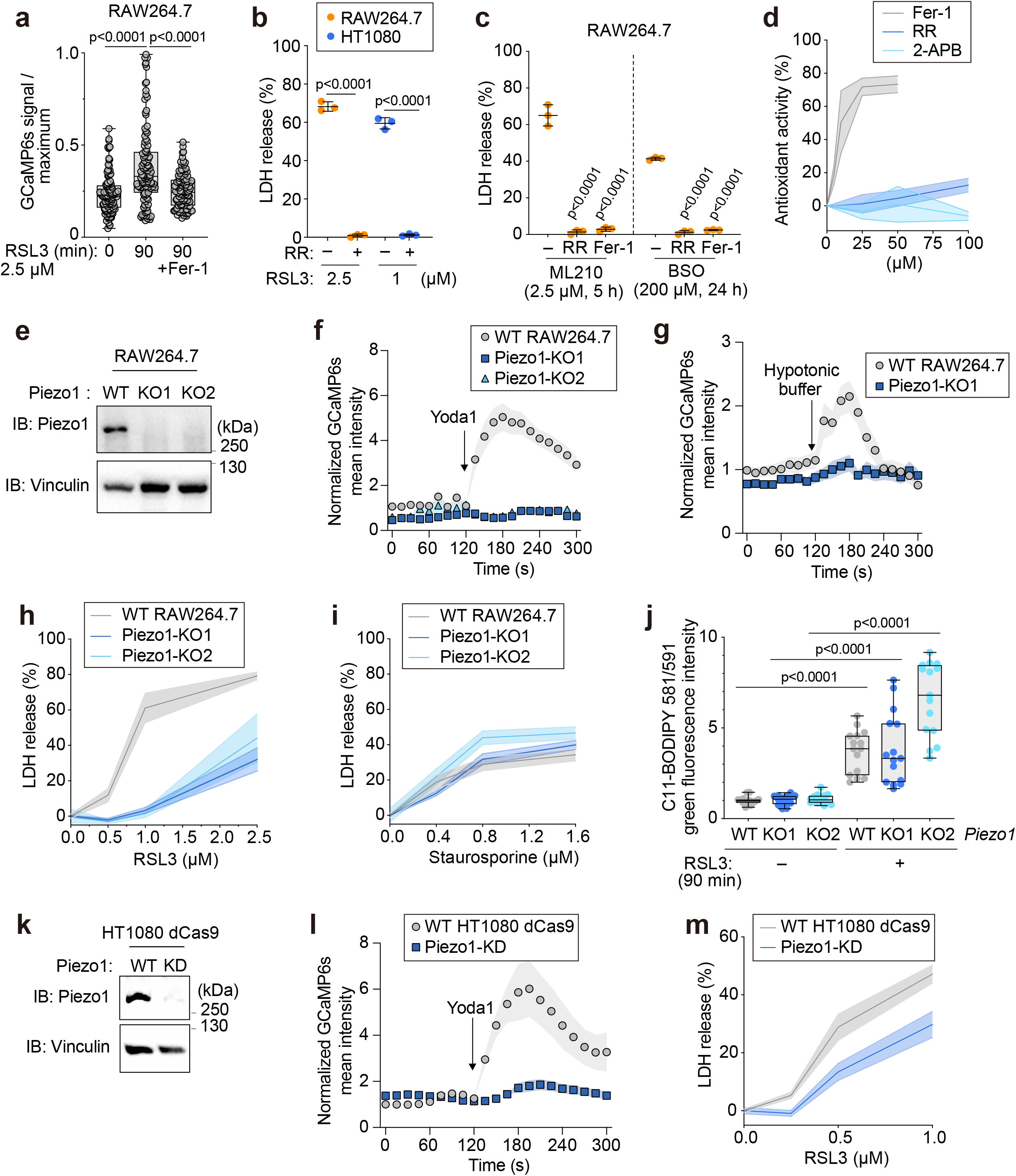
Piezo1 positively regulates ferroptosis. a) RAW264.7 cells expressing GCaMP6s +/-2.5 μM RSL3 +/-2.5 μM Fer-1 were video recorded for 5 min. Dots represent the average GCaMP6s signal over the maximum GCaMP6s fluorescence (ionomycin treated at the end of the recording, see Methods) per cell. Box plots are from 110-121 cells, n=3. (b) LDH release from RAW264.7 or HT1080 cells treated with RSL3 (2.5 μM and 1 μM, respectively) for 4 h +/-50 μM ruthenium red (RR) or 2.5 μM Fer-1. c) LDH release from RAW264.7 cells treated with 2.5 μM ML210 for 5 h or 200 μM BSO for 24 h +/-50 μM RR or 2.5 μM Fer-1. d) the antioxidant activity of RR, 2-APB and Fer-1 was assayed using DPPH. e) Immunoblot for Piezo1. f-g) Time-lapse GCaMP6s imaging of RAW264.7 cells treated with 10 μM Yoda1 (f) or exposed to hypotonic shock (∼100 mOsm) when indicated (g). i-h) LDH release from WT or Piezo1 KO cells treated with the indicated concentrations of RSL3 for 4 h (i) or staurosporine for 6 h (h). p=0.0015 (2.5 μM RSL3, WT vs Piezo1 KO1); p=0.0069 (2.5 μM RSL3, WT vs Piezo1 KO2); p=0.8343 (1.6 μM STS, WT vs Piezo1 KO1); p=0.0428 (1.6 μM STS, WT vs Piezo1 KO2). j) C11-BODIPY 581/591 imaging in WT and Piezo1 KO cells treated with 2.5 μM RSL3 for 90 min. n=3, 5 fields per n. k) Immunoblot for Piezo1. l) Time-lapse GCaMP6s imaging of HT1080 cells treated with 10 μM Yoda1. m) LDH release from WT and Piezo1 KD HT1080 cells treated with the indicated concentrations of RSL3 for 4 h. In all cases, data are shown as mean ± SD, n=3.

The preceding observations, together with the finding that plasma membrane tension increases during ferroptosis, prompted us to investigate the involvement of Piezo1. We edited *Piezo1* using CRISPR/Cas9 in RAW cells and established two independent Piezo1 KO lines (Fig. 4e); effective elimination of the channel was confirmed by the observation that the Ca^2+^ influx normally induced by Yoda1 (a Piezo1 agonist) in WT cells –determined following GCaMP6S transfection– was completely abrogated in the KO lines (Fig. 4f). We next tested whether increasing plasma membrane tension hydrostatically, i.e. by applying the same hypotonic shock as in Fig. 3f, similarly caused Ca^2+^ influx in a Piezo1-dependent manner. This was indeed the case: cytosolic [Ca^2+^] increased within seconds of hypotonic stress in WT but not in Piezo1-KO cells (Fig. 4g).

To test the role of Piezo1 in ferroptosis, we inhibited GPX4 using RSL3. Both Piezo1 KO cell lines were highly resistant to RSL3-induced cell death (Fig. 4h), but not to apoptosis induced by staurosporine (Fig. 4i). The deletion of Piezo1 did not affect the production of lipid peroxides, detected using C11-BODIPY 581/591 (Fig. 4j), suggesting that Piezo1 positively regulates ferroptotic cell death downstream of lipid peroxidation. To validate these results in a different experimental system, we generated a PIEZO1 stable knockdown (KD) line in dead Cas9 (dCas9) expressing HT1080 cells using a CRISPR interference (CRISPRi) system (Fig. 4k)^23, 24^. That Piezo1 was effectively knocked down was confirmed functionally in the cells by recording Yoda1-induced Ca^2+^ influx (Fig. 4l). As expected, knocking down the expression of PIEZO1 blunted their release of LDH in response to GPX4 inhibition as compared to their WT counterparts, further establishing a role of Piezo1 in promoting ferroptosis (Fig. 4m).

### Lipid peroxide accumulation collapses monovalent cation gradients via activation of Piezo1 and inhibition of the Na^+^/K^+^ ATPase

Na^+^ and K^+^ are the most abundant cations in the extracellular fluid and cytosol, respectively, thereby playing a major role in regulating cell volume^25^. The preceding experiments demonstrated that lipid peroxidation increased cell volume and plasma membrane tension, and that the deletion of Piezo1 protected cells from ferroptosis. We reasoned that lipid peroxidation activated monovalent cation fluxes (at least in part) via Piezo1, resulting ultimately in cell rupture. To directly test this assumption, we employed atomic absorption spectrometry (AAS) to determine the cellular concentrations of Na^+^ and K^+^ in RSL3-treated cells, calculated using the cell volumes that were measured in parallel. We found that the intracellular concentration of Na^+^ started to increase sharply ∼60 min after GPX4 inhibition, while the intracellular K^+^ decreased concomitantly (Fig. 5a). Within 90 min –before the cells became permeable to the small dye PI (Fig 1b)– the cytosolic cation concentrations approximated those of the extracellular fluid, suggesting that monovalent cation gradients were entirely dissipated before cell rupture. Importantly, these changes were suppressed in the presence of Fer-1 (Fig. 5b,c). A similar collapse of the cation gradients was observed in HeLa cells within 150 min (Fig. 5d), a time point when only 10% of the cells were permeable to PI (Fig. 1c). As an additional measure of the status of the Na^+^/K^+^ gradient, we determined if the membrane potential was affected during the induction of ferroptotic cell death using the potential-sensitive dye DiBAC_3_(4), an anionic fluorescent probe that accumulates in depolarized cells^26^. We found that inhibiting GPX4 depolarized RAW cells with a time course that approximated the K^+^ loss measured by AAS. In fact, after >60 min of GPX4 inhibition, the cells were depolarized to the same extent as were cells bathed in high K^+^ medium (Fig. 5e), a condition reported to cause nearly complete loss of the resting membrane potential in mammalian cells^27^. Taken together, these data supported the finding that lipid peroxidation is associated with a marked collapse of the monovalent cation gradients.

**Figure 5.**
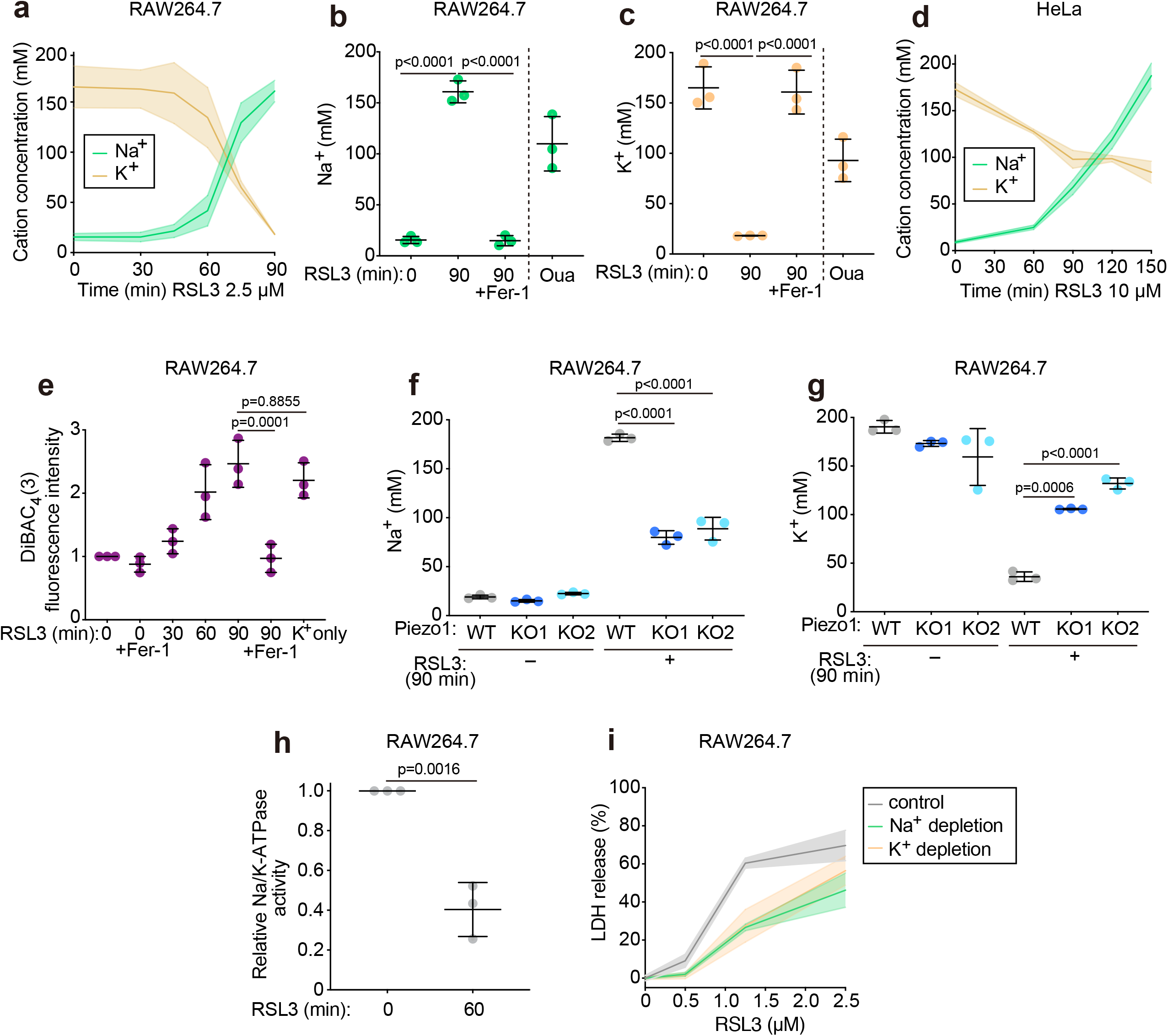
Lipid peroxide accumulation collapses monovalent cation gradients via the activation of Piezo1 and the inhibition of the Na^+^/K^+^-ATPase. a-d) [Na^+^] and [K^+^] determined by atomic absorption for cells treated with RSL3 over time. In (b) and (c), 2.5 μM Fer-1 was given at the time of RSL3 treatment or 1 mM ouabain (Oua) was given for 30 min. e) DiBAC_4_(3) imaging in RAW cells treated with 2.5 μM RSL3 for the indicated periods +/-2.5 μM Fer-1. f-g) [Na^+^] and [K^+^] were determined as in (a) for WT and Piezo1 KO cells. Data shown are the mean ± SD, n=3. h) Na^+^/K^+^ ATPase activity in RAW cells +/-2.5 μM RSL3 for 60 min. i) LDH release from cells ± 2.5 μM RSL3 for 3 h in control medium or medium in which N-methyl-D-glucamine^+^ (NMG^+^) or K^+^ was substituted for Na^+^. In all cases, data are shown as mean ± SD, n=3. p=0.0006 (1.25 μM RSL3, control vs NMG^+^ medium); p=0.0007 (1.25 μM RSL3, control vs K^+^ medium).

In order to discern the contribution of Piezo1 to the loss of cation gradients, we measured the intracellular concentrations of Na^+^ and K^+^ in Piezo1 KO cells by AAS. Remarkably, even after 90 min of GPX4 inhibition, the Piezo1 KO cells maintained their K^+^ gradient and the gain in cytosolic Na^+^ was strongly suppressed (Fig. 5f,g). This result would suggest that Piezo1 serves as a significant conduit for monovalent cation permeability in cells undergoing ferroptosis.

That the activation of Piezo1 alone, however, would suffice to collapse monovalent cation gradients would be surprising given the robust activity of the Na^+^/K^+^-ATPase, which continuously pumps 3 Na^+^ out of the cells in exchange for 2 K^+^ using the energy derived from the hydrolysis of a single ATP^28^. In otherwise untreated cells, inhibition of the Na^+^/K^+^-ATPase with ouabain for 30 min suffices to deplete K^+^ while causing a concomitant gain in Na^+^, yielding a phenotype similar to that observed following inhibition of GPX4 (Fig. 5b,c). We therefore analyzed whether ferroptosis is also associated with inhibition of the Na^+^/K^+^ pump. To this end, we measured the ouabain-sensitive fraction of the ATPase activity of a membrane-enriched preparation of cells, detecting phosphate released from ATP using malachite green. As illustrated in Figure 5h, the Na^+^/K^+^-ATPase activity of cells treated with RSL3 for 1 h was inhibited by ∼60% compared to untreated controls, suggestive of a contribution of Na^+^/K^+^-ATPase inhibition to the collapse of the monovalent cation gradients. Collectively, these data suggest that the loss of monovalent cation gradients observed during ferroptosis reflects the combined effects of inhibition of the Na^+^/K^+^-ATPase and the concomitant activation of Piezo1.

We further investigated whether the collapse in cation gradients and membrane potential are essential steps in the development of ferroptosis. This was accomplished by manipulating the ionic composition of the extracellular medium while the cells were exposed to the GPX4 inhibitor. To this end, we first substituted the external Na^+^ for N-methyl-D-glucamine (NMG^+^), a larger organic cation that cannot be transported by cation channels. This experiment was revealing: when curtailing Na^+^ influx, the cells were less susceptible to ferroptosis (Fig. 5i). Similar results were observed in cells incubated in K^+^-rich (instead of Na^+^-rich) medium (Fig. 5i), suggesting that Na^+^ gain and/or K^+^ loss sensitize the cells for the rapid cell death that accompanies GPX4 inhibition.

### Piezo1 and TRP channels cooperatively potentiate ferroptosis

The activation of Piezo1 accounted for approximately half of the change in intracellular Na^+^ and K^+^ induced by RSL3 (Fig. 5g,h). Accordingly, the cation fluxes were only partially suppressed in Piezo1-deficient cells, which were only partially resistant to RSL3. In contrast, cells treated with RR were completely refractory to ferroptosis (Fig. 4b,h,m). These observations suggested the involvement of other (mechanosensitive) cation channels, possibly members of the TRP family, which also enable the permeation of monovalent ions and are known to be blocked by RR. TRP channels are also inhibited with 2-APB ^29^. On treating RAW or HeLa cells with 2-APB, we found that the Na^+^ influx and K^+^ efflux induced by RSL3 were markedly suppressed (Fig. 6a,b and Supplementary Fig. 4a,b). While the ion fluxes were depressed to a level approaching that seen with Fer-1, blocking TRP channels with 2-APB did not affect the oxidation of lipids upon GPX4 inhibition (Fig. 6c), in good accordance with DPPH antioxidant assays where 2-APB showed no antioxidant activity (Fig. 4c). LDH release assays were performed to determine whether preventing the increase in cation permeability with 2-APB impacted the onset of lytic cell death. As documented in Figure 6d-e and Supplementary Fig. 4c, 2-APB strongly suppressed RSL3-induced death in RAW, HeLa, and HT1080 cells. In addition to blocking TRP channels, 2-APB also inhibits the inositol 1,4,5-*tris*phosphate receptor (IP3R), an ER-resident Ca^2+^ channel^30^. To determine whether the latter target contributed to the anti-ferroptotic effect of 2-APB, we also tested Xestospongin C, a selective IP3R inhibitor^31^. In contrast to 2-APB, Xestospongin C did not affect RSL3-induced cell death (Fig. 6d), supporting the importance of TRP channels in ferroptosis.

**Figure 6.**
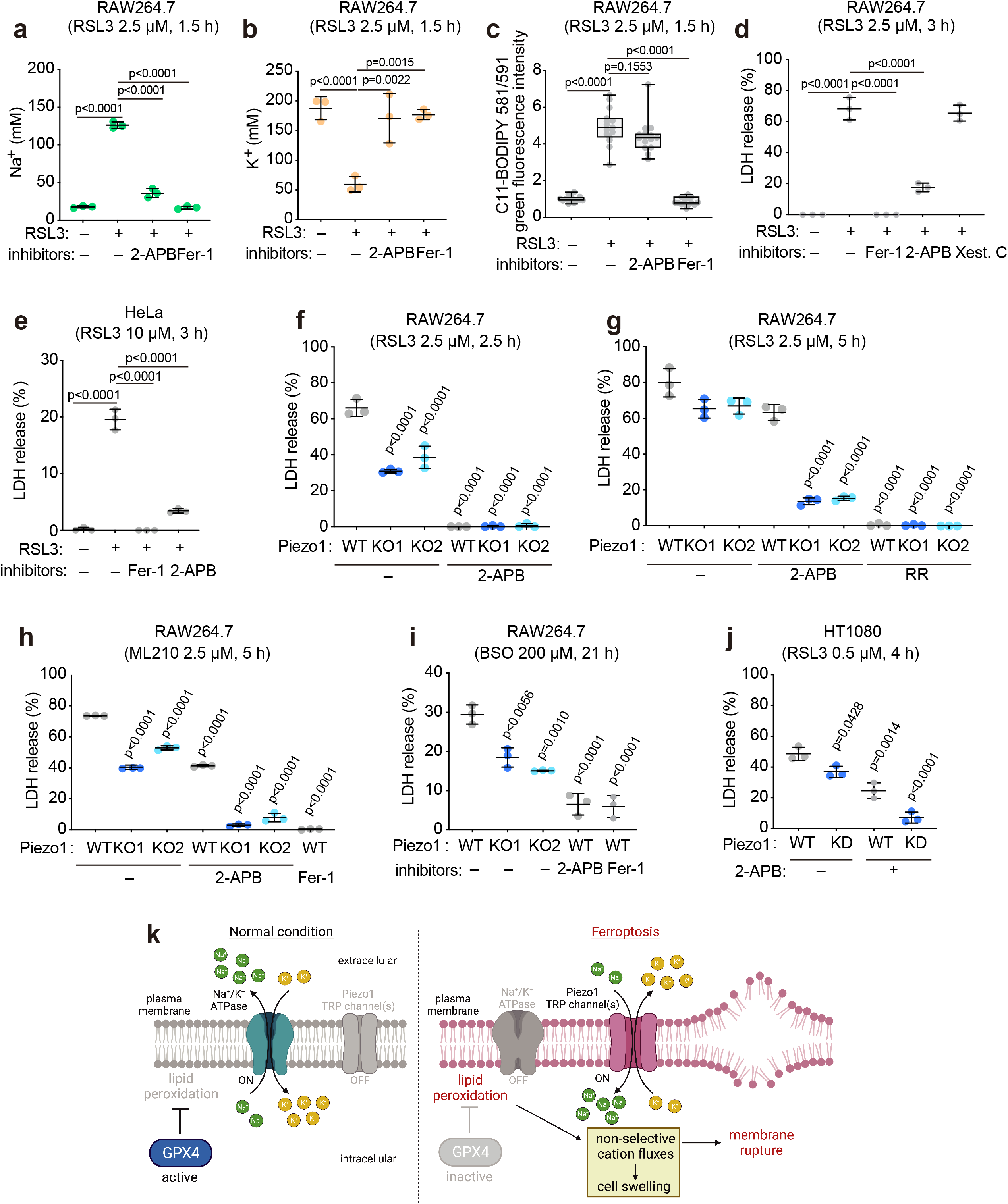
Piezo1 and TRP channels cooperatively potentiate ferroptosis. a-b) [Na^+^] and [K^+^] determined by atomic absorption for cells treated with RSL3 ± 50 μM 2-APB or 2.5 μM Fer-1. c) Determinations of the green (oxidized) C11-BODIPY 581/591 signal in cells treated as in (a-b). d) LDH release from cells treated with 2.5 μM RSL3 for 3 h ± 2.5 μM Fer-1, 50 μM 2-APB or 10 μM Xestospongin C (Xest C). e-f) LDH release from WT and Piezo1 KO cells treated with 2.5 μM RSL3 for 2.5 h (e) or 5 h (f) ± 50 μM 2-APB or 50 μM ruthenium red (RR). g-h) LDH release from WT or Piezo1 KO cells ± 2.5 μM ML210 for 5 h (g) or 200 μM BSO for 21 h (h) ± 50 μM 2-APB or 2.5 μM Fer-1. i) LDH release from WT or Piezo1 KD HT1080 cells treated with 0.5 μM RSL3 for 4 h ± 50 μM 2-APB. In all cases, data are shown as the mean ± SD, n=3. j) Schematic representation of monovalent cation transport pathways at rest and during ferroptosis.

To assess the potentially cooperative role of Piezo1 and TRP channels in ferroptosis, we treated WT or Piezo1 KO cells with 2-APB, allowing lipid peroxides to accumulate for longer time periods than detailed in Figure 5. After 2.5 h of GPX4 inhibition, Piezo1 KO cells released ∼50% less LDH, whereas 2-APB almost completely blocked LDH release from WT and KO cells (Fig. 6e). After 5 h, 2-APB only partially suppressed LDH release in WT cells, but the combination of 2-APB and the deletion of Piezo1 almost fully prevented it (Fig. 6f). This suggests that Piezo1 and TRP channels play a cooperative role in ferroptosis and that TRP channels may feature more prominently in its earlier stages. Similar results were obtained upon treatment with other ferroptosis inducers, i.e. ML210 and BSO (Fig. 6h,i), and in Piezo1 KD HT1080 cells (Fig. 6j).

In sum, these results support the model depicted in Figure 6k where oxidized plasma membrane lipids increase membrane tension, activate Piezo1 and TRP channels, and in parallel incapacitate the Na^+^/K^+^ ATPase. These effects collectively collapse monovalent cation gradients, cause cell volume gain and potentiate plasma membrane rupture at the onset of ferroptosis.

## Discussion

In this study, we established that the lipid peroxidation that occurs during ferroptosis takes place preferentially at the plasma membrane. It is noteworthy that phospholipid oxidation is reported to occur in various other subcellular locations, including mitochondria, the endoplasmic reticulum, lysosomes, lipid droplets, and peroxisomes, to an extent that varies depending on the cell type and the microenvironment^32^. It was nevertheless unclear how intracellular oxidized phospholipids coordinately lead to ferroptotic cell lysis. It was previously shown that the inhibition of GPX4 with RSL3 resulted in cell death even in cells depleted of mitochondria^5, 33^. While mitochondria are a major source of ROS, they are seemingly not themselves an essential target of the oxidation that causes ferroptosis. Two recent independent reports employing genetic screens support the notion that the plasmalemma is an important contributor. First, the ferroptosis suppressor protein 1 (FSP1) was recently identified for its ability to inhibit lethal peroxidation and ferroptosis. FSP1 is a lipophilic radical-trapping antioxidant that reduces ubiquinone to ubiquinol and vitamin K to its hydroquinone; the reduced products are potent antioxidants that trap oxygen radicals and prevents lipid peroxidation. Importantly, the ability of FSP1 to antagonize ferroptosis relies on its N-myristoylation, which targets it to the plasma membrane^34-36^. Secondly, the addition of exogenous monounsaturated fatty acids was found to confer resistance of cells to ferroptotic cell death^37^. These fatty acids incorporate in the plasmalemmal phospholipids but lack oxidizable polyunsaturated acyl chains, making the cell less susceptible to ferroptosis.

We assessed the loss of monovalent cation gradients across the plasma membrane in two ways: directly, via AAS to determine absolute cellular [Na^+^] and [K^+^], and indirectly by measuring the membrane potential. It is noteworthy that, at the time when the plasma membrane had depolarized (e.g. 60 min), no significant change in mitochondrial membrane potential (MMP) was detected (Fig. S2). Instead, we noted that MMP –an indicator of mitochondrial functional integrity– only became affected after 90 min, i.e. when cells begin to rupture. Depolarization of the MMP possibly resulted from Ca^2+^ influx (Fig. 3a), which is known to uncouple mitochondria and reduce MMP effectively^38^. Mitochondrial dysfunction therefore appeared to be the consequence and not the cause of ferroptosis.

As mentioned above, ferroptosis-associated lipid oxidation has been detected not only in mitochondria but also in lysosomes. It is therefore conceivable that oxidation of the phospholipids that form the limiting membrane of lysosomes may cause their rupture, which would release their resident hydrolases into the cytosol and initiate the death process. Such ectopic proteolysis could conceivably also cause the observed increase in plasmalemmal permeability. However, we also investigated lysosomal integrity, determined by their ability to retain H^+^, and did not observe any effects even after 90 min of GPX4 inhibition (Fig. S3), long after the changes in plasma membrane permeability were manifest.

Having ruled out mitochondrial and lysosomal damage as initiators of the ferroptotic events described here, we considered the role of cytosolic ions in the process. Gross changes in the ionic composition of the cells can be expected to be associated with alterations in cell volume. Accordingly, volume changes had been suggested to occur in cells undergoing ferroptosis^7, 14^; these suggestions, however, were based on indirect measurements and inconsistent with each other. The rounded shape adopted by cells undergoing ferroptosis had been interpreted to reflect cell volume shrinkage, without taking into account possible changes in cell height. On the other hand, cell swelling was proposed to occur based on observations using the nuclear tension sensor, cytosolic phospholipase 2 (cPLA2)-mKate, which purportedly translocates to the nucleus upon cell swelling^39, 40^. Interestingly, the translocation of cPLA2 to the nuclear envelope requires not only an increase in nuclear membrane tension, but also of cytosolic [Ca^2+^] ^9^; both of these events are expected to occur based on our findings. We measured changes in cell volume directly during GPX4 inhibition using electronic cell sizing and confirmed that the cells indeed underwent swelling, which we attributed to an imbalance in ion fluxes across the plasma membrane. Imaging of adherent cells confirmed that cell height and volume increased during ferroptosis.

Intriguingly, recent *in silico* simulations of molecular dynamics in membrane bilayers have revealed that peroxidation of phospholipids (e.g. during ferroptosis) can significantly change the physical properties of the membrane, causing its thinning and increased curvature^17^. Our findings demonstrating elevated plasma membrane tension upon lipid peroxidation support these models. Monovalent ion fluxes and cell death, but not lipid peroxide accumulation, were also strongly prevented by blocking or editing mechanosensitive channels. These results implicate membrane tension and the relative expression of mechanosensitive channels as major determinants in ferroptosis.

Members of the TRP family are generally non-selective cation channels and their opening probability is directly affected by changes in physical properties of the membrane such as tension, thickness, curvature, and lipid order ^41^. TRPC3 and TRPC6 are expressed in sensory neurons and cochlear hair cells and have been shown to be required for their normal mechanotransduction^42^. TRPM7 is activated in response to membrane stretch by osmotic swelling, and induces cation fluxes, thereby contributing to cell volume regulation^43^. It is reasonable to assume that TRP channels directly sense phospholipid peroxidation-induced changes in membrane properties, becoming activated to facilitate non-selective cation efflux/influx. Alternatively, products from hydroperoxidized phospholipids, i.e. peroxidized fatty acids like hydroperoxyeicosatetraenoic acids (HpETEs) and hydroxyoctadecadienoic acids (HODEs), or their reactive aldehyde metabolites, including 4-hydroxy-2-nonenal (4-HNE) and malondialdehyde (MDA), could activate TRP channels during ferroptosis as an autocrine ligand or through oxidative modifications^44, 45^. It has been shown that 12(*S*)-HpETE and 15(*S*)-HpETE are potent activators for TRPV1, whereas 9(*S*)-HODEa and 13(*S*)-HODE activate TRPV1 and TRPA1, possibly by serving as ligands^45^. Meanwhile, highly reactive aldehydes including 4-HNE have been shown to induce covalent modifications of several amino acid residues of TRPV1 (Cys621) and TRPV (Cys619, Cys639, Cys663 and Lys708)^45^.

This report is also the first to directly determine the involvement of Piezo1 in ferroptosis. Piezo1 is directly activated by increased membrane tension and is only slightly (2-fold) more permeable to Ca^2+^ than to Na^+^ and K^+^ ^20, 46^. Our genetic and pharmacological approaches showed that Piezo1 contributes to the Na^+^ gain, K^+^ loss, and cell rupture associated with ferroptosis. Thus, as phospholipids become oxidized and the permeability of the membrane to monovalent ions increases, Piezo1 jointly with TRP channels initiate a feed-forward mechanism: as they contribute to cell swelling, the channels become further activated, increasing plasma membrane tension and self-gating. Though not definitive, our findings suggest that TRP channels may be early initiators of ferroptosis while Piezo1 may accelerate the death process.

We also found that the Na^+^/K^+^ ATPase is less efficient in cells undergoing ferroptosis. The peroxidation of the surrounding phospholipids may alter the bilayer in a manner that diminishes the activity of the pump. Crystal structures of the Na^+^/K^+^ ATPase have revealed three binding pockets for cholesterol and phospholipids^47-50^. Subsequent *in vitro* studies using liposome reconstitution systems suggested that each one of the binding pockets has a different affinity for lipid species and exerts distinct effects on the enzymatic activity: PS/cholesterol are stabilizing, PE is stimulatory, and PC or sphingomyelin/cholesterol are inhibitory^51, 52^. Interestingly, the stimulatory effect of PE was maximized when the *sn*-1/2 acyl moieties were C18:0/C20:4 or C18:0/C22:6 ^51, 52^. Given the fact that ferroptosis is triggered by preferential peroxidation of polyunsaturated fatty acids, notably C20:4 (arachidonic acid) and C22:6 (docosahexaenoic acid)^53^, we speculate that peroxidization of PE may reduce its stimulatory effect on Na^+^/K^+^ ATPase activity. To date there is no evidence that specific Na^+^/K^+^ ATPase residues are modified by reactive aldehydes, though early studies have revealed their association with reduced pump activity^54-56^. Intriguingly, the Na^+^/K^+^ ATPase activity is known to be reduced by S-glutathionylation of the cysteine residues of the α1 subunit (Cys244, Cys454, Cys458, and Cys459), and of the β1 subunit (Cys46) ^44^. These cysteine residues might be potential target sites for reactive aldehydes to exert a negative effect on Na^+^/K^+^ ATPase activity.

There is precedent for a role of monovalent cation flux in programmed cell death in both apoptosis and pyroptosis. It has been shown that K^+^ efflux during apoptosis is triggered by multiple K^+^ channels, including voltage-gated K^+^ channels and Ca^2+^-activated K^+^ channels, depending on the cell types and stimuli, which consequently induces cell shrinkage^57^. During apoptosis, Na^+^ influx also occurs and is thought to be mediated by multiple types of Na^+^ channels, such as voltage-dependent and -independent channels (i.e. epithelial Na^+^ channels), which have been shown to augment PS exposure^57^. In pyroptosis, it is now well-established that K^+^ efflux is the major trigger of inflammasome formation, a process mediated by low K^+^-sensing effectors (e.g. NIMA-related kinase 7) that activate NOD-like receptor protein 3 (NLRP3)^58^. K^+^ efflux leading to pyroptosis can be induced by diverse means including by K^+^ ionophores (e.g. nigericin), pore-forming toxins, pannexin channels, etc. and is accelerated by gasdermin D and NINJ1.

Here, we have established that ferroptosis is also associated with cation gradient collapse via the activation of channels, but not the oligomeric NINJ1 complex, proposed to form a pore. While much remains to be understood about the mechanisms that ultimately lead to ferroptotic cell death, our observations implicate the role of biophysical parameters of the plasma membrane upon its oxidation, hydrostatic pressure, and the expression/gating of Piezo1 and TRP channels as orchestrators of the entire process.

## Acknowledgments

We thank Dr Nobuhiko Kayagaki and Dr Vishva Dixit for providing the rabbit monoclonal anti-mouse NINJ1 antibody. This work was supported by JSPS/MEXT KAKENHI Grant Numbers JP20K07011, JP20KK0361 (Y.H.); JP21H02620 and JP21H00268 (A.M.) and by the Takeda Science Foundation. S.A.F. is funded by a CIHR Project Grant PJT169180. S.G. was supported by grant FDN-143202 from CIHR.

## Materials and Methods

### Reagents

1*S*, 3*R*-RSL3 (catalog #SML2234), L-buthionine sulfoximine (BSO, catalog #B2515), ML210 (catalog #SML0521), ferrostatin-1 (catalog #SML0583), propidium iodide (PI; catalog #P4170), H_2_O_2_ (catalog #H1009), staurosporine (catalog #S5921), lipopolysaccharide (catalog #L9764), valinomycin (catalog #V0627), cresyl violet (catalog #C5042), ionomycin (catalog #407950), ouabain (catalog #O3125), and Rhodamine-123 (catalog #R8004) were purchased from Sigma-Aldrich (Burlington, MA, USA). Deferoxamine mesylate (catalog #D9533) and xestospongin C (catalog #64950) were purchased from Cayman (Ann Arbor, MI, USA). BODIPY 581/591 C11 (catalog #D3861), DiBAC_4_(3) (catalog #B438), 10 kDa Alexa 647-dextran (catalog #D22914) and the protease inhibitor cocktail (catalog #A32955) were from Thermo Fisher Scientific (Waltham, MA, USA). Yoda 1 (catalog #5586) and 2-aminoethoxydiphenyl borate (catalog #1224) were from Tocris Bioscience (Bristol, UK). Concanamycin A (catalog #ab144227) and nigericin (catalog #ab120494) were from Abcam (Cambridge, UK). Antibodies used for immunoblotting were as follows: anti-Piezo1 (Proteintech, catalog #15939-1-AP); anti-Vinculin (Sigma, catalog #MAB3574); anti-GAPDH (Santa Cruz, catalog #sc-25778). Rabbit monoclonal anti-mouse NINJ1 antibodies were generously provided by Drs. Nobuhiko Kayagaki and Vishva Dixit (Genentech Inc, San Francisco). All antibodies were used at 1:1000 dilution for immunoblotting.

### Cell culture

HT1080 and HeLa cells were cultured in Dulbecco’s Modified Eagle’s medium (catalog #319-007-CL) containing 10% heat-inactivated fetal bovine serum in 5% CO_2_ at 37°C. RAW264.7 cells were cultured in RPMI 1640 medium (catalog #350-000-CL; Wisent, Saint-Jean-Baptiste, Canada), also containing 10% heat-inactivated fetal bovine serum in 5% CO_2_ at 37°C. In Fig 5c and 5d, RPMI 1640 medium was replaced by the following *isotonic media* just before RSL3 treatment: *normal medium*, 20 mM HEPES [pH 7.4], 150 mM NaCl, 5 mM KCl, 2 mM CaCl_2_, 1 mM MgCl_2_, 10 mM glucose); *Na*^*+*^*-depleted medium*, where NaCl 150 mM in the normal medium was substituted with 150 mM N-methyl-D-glucamine (NMG)-Cl; *K*^*+*^ *high medium*, where the concentrations for NaCl and KCl in the normal medium were swapped (NaCl 5 mM and KCl 150 mM). To induce ferroptosis with BSO, cells were treated with BSO in the presence of 100 μM Fe(NH4)_2_(SO4)_2_ (Fig. 1k, 4c, and 6i).

### Gene editing

RAW-ASC cells (catalog #raw-asc, Invivogen, San Diego, CA, USA), RAW264.7 cells stably expressing apoptosis-associated speck-like protein containing a CARD (ASC), were used as parental cells. RAW-ASC cells were transfected with custom CRISPR gRNA plasmid DNA (U6-gRNA:CMV-Cas-9-2A-tGFP) (catalog #CAS9GFFP, Sigma-Aldrich) using the FuGENE HD transfection reagent (catalog #E2311, Promega, Madison, WI, USA). The NINJ1 target region sequence was GCCAACAAGAAGAGCGCTG. To generate RAW264.7 Piezo1 KO cells, the CRISPR gRNA plasmid encoding the GFP, Cas9 and predesigned gRNA (CAGCCCGAAGATGCTTATG) was purchased from Sigma-Aldrich. In both cases, 24 h after transfection the cells were FACS-sorted for GFP into 96 well plates. Individual colonies were expanded and tested for NINJ1 or Piezo1 expression by immunoblotting.

For Piezo-KD HT1080 cells, parental HT1080 cells expressing stable nuclease-dead Cas9 (dCas9-BFP-KRAB, Addgene # 46911) were used as described previously^23^. The sgRNA targeting Piezo1 was designed previously^59^ and synthesized by Thermo Fisher Scientific. To knockdown Piezo1 in HT1080 dCas9 cells, the specific sgRNA was subcloned using BstXI and BlpI restriction enzymes into pU6-sgRN EF1Alpha-puro-T2A-BFP vector (Addgene #60955)^24^. A stable population of the HT1080 dCas9 cells expressing the plasmid containing the sgRNA were obtained by puromycin selection (3 μg/ml). Piezo1 deletion was confirmed in clones by immunoblotting.

### Microscopy

Cells were seeded onto 18-mm circular coverslips in 12-well plates. For image acquisitions, coverslips were mounted in a Chamlide CMB chamber (Quorum Technologies, Puslinch, Canada). Images were acquired using a spinning-disc confocal microscope (WaveFX; Quorum Technologies) operated by Volocity software (Quorum Technologies) as described previously^59^. Bright-field images (Fig. 2a and 2b) and fluorescence images for PI staining (Fig. 1b and 1c) were obtained using the EVOS M5000 Imaging System (Thermo Fisher Scientific).

### Cell death assays

Lactate dehydrogenase (LDH) assays were performed using the Cytotoxicity LDH Assay Kit (catalog #CK12, Dojindo, Kumamoto, Japan) according to the manufacturer’s protocol. The activity level of the LDH released into the culture media is presented as a percentage of the total activity level of LDH, as described previously^60^. For PI staining, cells were treated with 5 μg/mL PI for 15 min, observed under a microscope, and quantified as the percentage of PI-positive cells using ImageJ.

### Lipid ROS imaging

RAW264.7, HeLa, and HT1080 cells were seeded on glass coverslips. After stimulation, cells were treated with 5 μM BODIPY 581/591 C11 for 15-30 min before observation by spinning-disc confocal microscopy. Oxidized lipid levels were quantified using Volocity software (Quorum Technologies) and defined by the green fluorescence intensity, after background subtraction.

### Relative membrane potential determinations

RAW264.7 cells were seeded on glass coverslips. RPMI 1640 Medium was changed to the normal medium just before RSL3 treatment. Each coverslip was mounted in a chamber preincubated with the same medium containing 1 μM DiBAC_4_(3) for 5 min to make its binding to the chamber reach equilibrium, and cells on the coverslips were treated with 1 μM DiBAC_4_(3) for 10 min before microscopic observation. To depolarize the cells, they were treated with K^+^ high medium for 30 min. Average green fluorescence intensity in 5 fields for each sample was quantified using Volocity software, and is presented as relative fluorescence intensity (mean ± SD from 3 independent experiments).

### Mitochondrial membrane potential measurement

RAW264.7 cells were seeded on glass coverslips and treated with the indicated reagents. 5 μM Rhodamine-123 was added 15 min before observation by fluorescence microscopy. Average green fluorescence intensity in 3 fields for each sample was quantified by Volocity software and is presented as relative fluorescence intensity (mean ± SD from 3 independent experiments).

### Calcium imaging

To measure the changes in intracellular (cytosolic) Ca^2+^ concentration [Ca^2+^]_i_, we utilized the fluorescent Ca^2+^-sensing protein GCaMP6s that has a much greater dynamic range and faster response kinetics than other commonly used probes^61^. In Fig. 4a, to express GCaMP6s, RAW264.7 cells seeded onto coverslips were transfected with GCpGP-CMV-GCaMP6s (Addgene plasmid 40753) using FuGENE HD (Promega), and treated with or without RSL3 and Fer-1, as indicated. Coverslips were mounted in chambers with Tyrode’s buffer (10 mM HEPES [pH 7.4], 140 mM NaCl, 10 mM glucose, 5 mM KCl, 2 mM CaCl_2_, 1 mM MgCl_2_), and then cells (3 separate fields per sample) were subsequently imaged in Tyrode’s buffer every 15 sec for 5 min. Two min after starting recording, cells were treated with 5 μM ionomycin to induce a saturating increase in [Ca^2+^]_i_ for 3 min. Using ImageJ to analyze the video recordings, regions of interest were generated to determine the average fluorescence signal before (S1), and after ionomycin treatment to determine the maximum fluorescence signal (S2). In order to compare [Ca^2+^]_i_ between cells with different expression levels of GCaMP6s, the data were normalized as fluorescence ratios (S1/S2, where a high ratio indicates high [Ca^2+^]_i_); the normalized values are shown as a violin plot graph (n=110-121, from 3 independent experiments). RAW cells expressing GCaMP6s-GFP were incubated with Tyrode’s buffer for 10 min before imaging. Recordings were made every 15 s for 5 min and 10 μM Yoda1 was added at the 2 min time point. Data are presented as the mean ± SEM, >15 cells, n=3.

### Lysosomal integrity assessment

To assess the integrity of lysosomes, we performed cresyl violet staining as described previously^62^ with minor modifications, enabling visualization of acidic compartments within the cells. Briefly, RAW264.7 cells were seeded onto coverslips and treated with 5 μM Alexa 647-dextran 10 kDa overnight. The next day, medium was refreshed and then cells were incubated at 37°C for 1 h; this chase period led to selective retention of Alexa 647-dextran 10 kDa in lysosomes. Cells were treated with 1 μM cresyl violet for 1 min, washed with PBS, and subjected to microscopic observation. Average red and far-red fluorescence intensity in 3 fields for each sample was quantified using Volocity, and is presented as relative fluorescence ratio, where the red fluorescence intensity of a field was divided by the far-red intensity (cytosol background was subtracted, mean ± SD from 3 independent experiments).

### Fluorescence lifetime imaging microscopy (FLIM)

Cells were incubated with 2□μM Flipper-TR (Spirochrome, Catalog #SC020) for 5□min in PBS before imaging. Frequency-domain FLIM measurements were collected using a Zeiss AxioObserver inverted microscope equipped with a Lambert-FLIM attachment, using a ×40/1.4 NA oil immersion objective and an Orca R2 CCD camera. The frequency of modulation was 400□MHz and the instrument was calibrated based on the fluorescence lifetime of lucifer yellow CH, lithium salt (Molecular Probes, Catalog #L453) assuming a monoexponential lifetime of 5.7 ns^63^. Lifetime analysis was performed using FLIM software from Lambert Instruments.

### Antioxidant activity assay

Antioxidant activity was assayed using DPPH (Dojindo, catalog #D678) according to the manufacturer’s protocol with minor modifications. In brief, each (putative) antioxidant was dissolved in 10 μL ethanol at different concentrations, and mixed with 40 μL of the assay buffer and 50 μL of the DPPH working solution. After incubation at room temperature for 30 min, absorbance at 517 nm was measured using a microplate reader.

### Electronic cell sizing

Cell volume measurements were performed as previously described with minor modifications^64^. Briefly, RAW264.7 cells and HeLa cells were lifted mechanically or by trypsinization, respectively. The mean cell diameter for >10,000 cells was determined using the Z2 Coulter particle count and size analyzer (Beckman Coulter, Brea, CA, USA) and converted to volume by assuming that the suspended cells were spherical.

### Atomic absorption spectrometry

Atomic absorption spectrometry was performed as previously described with minor modifications^64^. In brief, RAW264.7 cells and HeLa cells were seeded onto 6-well plastic tissue culture plates. After stimulation, plates were washed x3 by gentle, brief submersion of the dishes in cold Na^+^/K^+^-free buffer (150 mM LiCl, pH 7.4) to remove contaminating ions. Cells were lysed with 1 mL H_2_O containing 1 mM HNO_3_. Lysis was done by scraping and vigorous pipetting. Samples were analyzed using a PinAAcle 900T Atomic Absorption Spectrometer (PerkinElmer, Waltham, MA, USA) to assess the total amount of ions. In parallel, cell numbers and cell volumes were determined using Z2 Coulter particle count and size analyzer (Beckman Coulter) to calculate total cell volume for each sample. Intracellular concentrations of Na^+^ and K^+^ were calculated by dividing the total ion content by the total cell volume.

### *In vitro* Na/K ATPase assay

RAW264.7 cells were seeded (3 × 10^6^ cells per 10 cm dish), treated with or without RSL3, and collected with ice-cold homogenizing buffer (50 mM HEPES [pH 7.4], 250 mM sucrose, 1 mM EGTA, protease inhibitor cocktail). Protein was extracted by Dounce homogenization with 50 strokes, and samples were centrifuged at low speed (3000 × *g*, 10 min, 4°C). Then, the supernatant was collected and ultracentrifuged (100,000× *g*, 30 min, 4°C). The resultant pellet was resuspended with 50 μL homogenizing buffer, and analyzed for ATPase activity in reaction buffer (50 mM HEPES [pH 7.4], 50 mM NaCl, 25 mM KCl, 5 mM MgCl_2_, 5 mM ATP) with or without 2 mM ouabain at 37°C for 30 min while shaking. The reaction was stopped by adding 10% trichloroacetic acid, and samples were centrifuged to remove precipitated material (14,000 rpm, 5 min). Enzyme activity was assessed by measuring the amount of inorganic phosphate ([Pi]) produced using a malachite assay kit (catalog #K1501, Echelon, Salt Lake City, USA), and protein concentration was determined by a bicinchoninic assay kit (catalog #23225, Thermo Fisher Scientific). To calculate Na/K ATPase (ouabain-sensitive) enzyme activity, the following equation was used: ([Pi] without Ouabain – [Pi] with ouabain])/amount of protein. Data are shown as relative enzyme activity (mean ± SD from 3 independent experiments).

### Image analysis and statistics

Image handling, quantification, and analysis of fluorescence images were performed using either Volocity 6.3 software (Quorum Technologies) or ImageJ software (Fiji v. 2.3.0/1.53c). All the values are expressed as means ± SD, and statistical analyses were performed using GraphPad Prism software (v.9.3.0). All experiments were repeated at least three independent times. Two groups were compared using two-tailed Student’s t-test. Multiple-group comparisons were conducted using either the one-way ANOVA analysis or two-way ANOVA analysis followed by Tukey-Kramer or Dunnett’s test.

## Figure Legends

**Supplementary Figure 1. GPX4 inhibition causes accumulation of lipid peroxides in the plasma membrane in HeLa and HT1080 cells**. a-b) C11-BODIPY 581/591 imaging in HeLa (a) and HT1080 (b) cells treated with 20 μM or 1 μM RSL3, respectively, for the indicated periods ± 2.5 μM Fer-1. c) C11-BODIPY 581/591 imaging in RAW264.7 cells treated with ML210 or BSO as indicated. Scale bar, 10 μm.

**Supplementary Figure 2. RSL3 treatment leads to a small reduction in the mitochondrial membrane potential**. a-b) Rhodamine-123 imaging in RAW264.7 cells treated with either 2.5 μM RSL3 ± 2.5 μM Fer-1, 1 mM ouabain (Oua) for 60 min, or 1 μM valinomycin (Val) for 15 min. Representative images (a) and quantification of the green fluorescence intensity (b, mean ± SD, n=3) are shown. Scale bar, 10 μm.

**Supplementary Figure 3. Lysosomal integrity is not affected by RSL3 treatment**. a-b) Imaging of cresyl violet, and indicator of acidic pH, and 10 kDa Alexa 647-dextran in RAW cells treated with 2.5 μM RSL3 for the indicated periods or 1 μM concanamycin A (Con A) for 1 h. Representative images (a) and fluorescence ratio of red (cresyl violet) to far-red (dextran 10 kDa) (b, mean ± SD, n=3) are shown. Scale bar, 10 μm.

**Supplementary Figure 4. 2-APB prevents cation fluxes and cell death upon GPX4 inhibition in HeLa cells**. a-b) [Na^+^] and [K^+^] determined by atomic absorption for HeLa cells treated with RSL3 at 10 μM for 3 h ± 50 μM 2-APB or 2.5 μM Fer-1. c) LDH release from HeLa cells treated with 10 μM RSL3 for 3 h ± 50 μM 2-APB or 2.5 μM Fer-1. In all cases, data are shown as mean ± SD (n=3).

