## Supplementary figures and images for "Lipid peroxidation increases membrane tension, Piezo1 gating and cation permeability to execute ferroptosis"

### Supplemental Figures

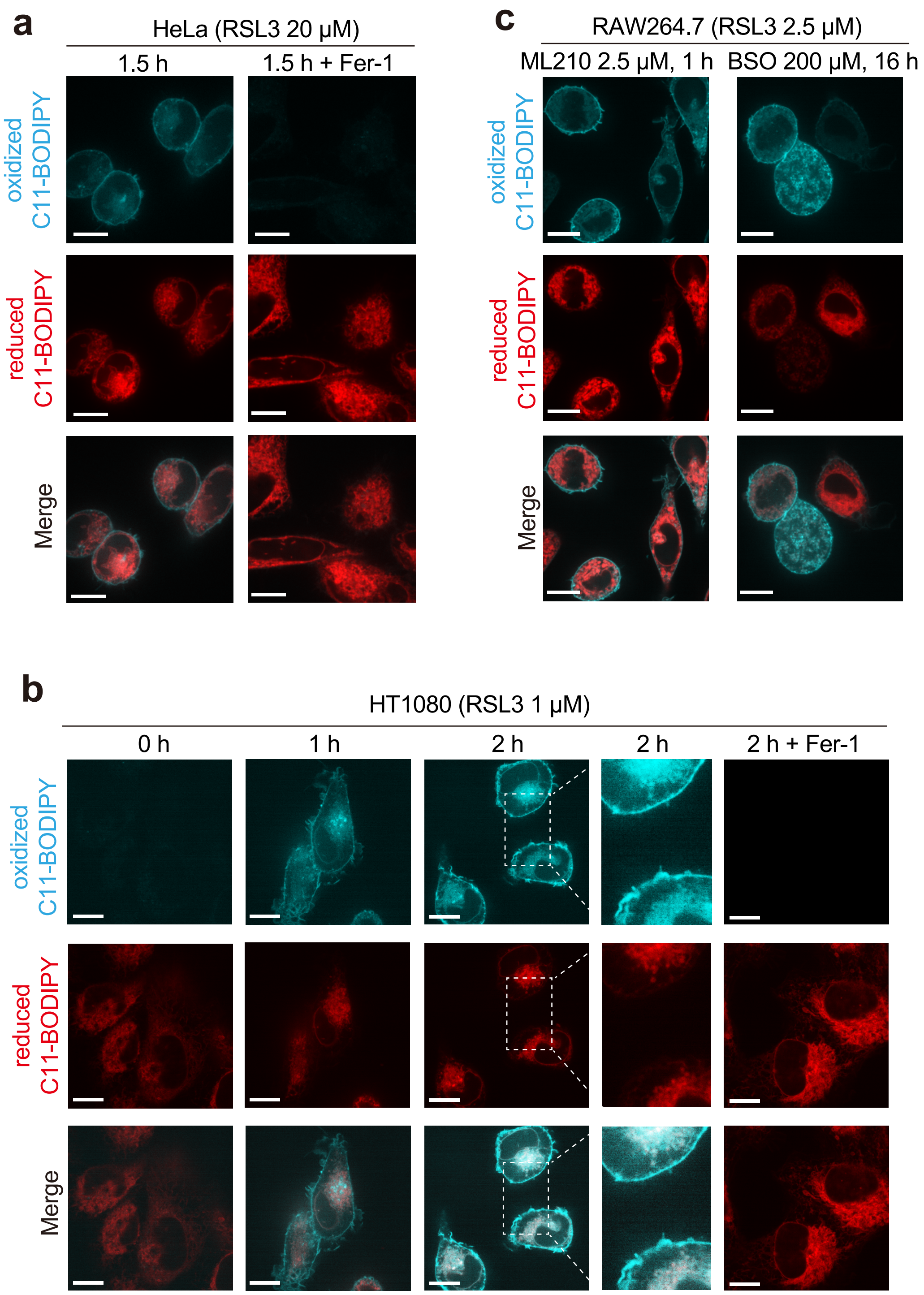

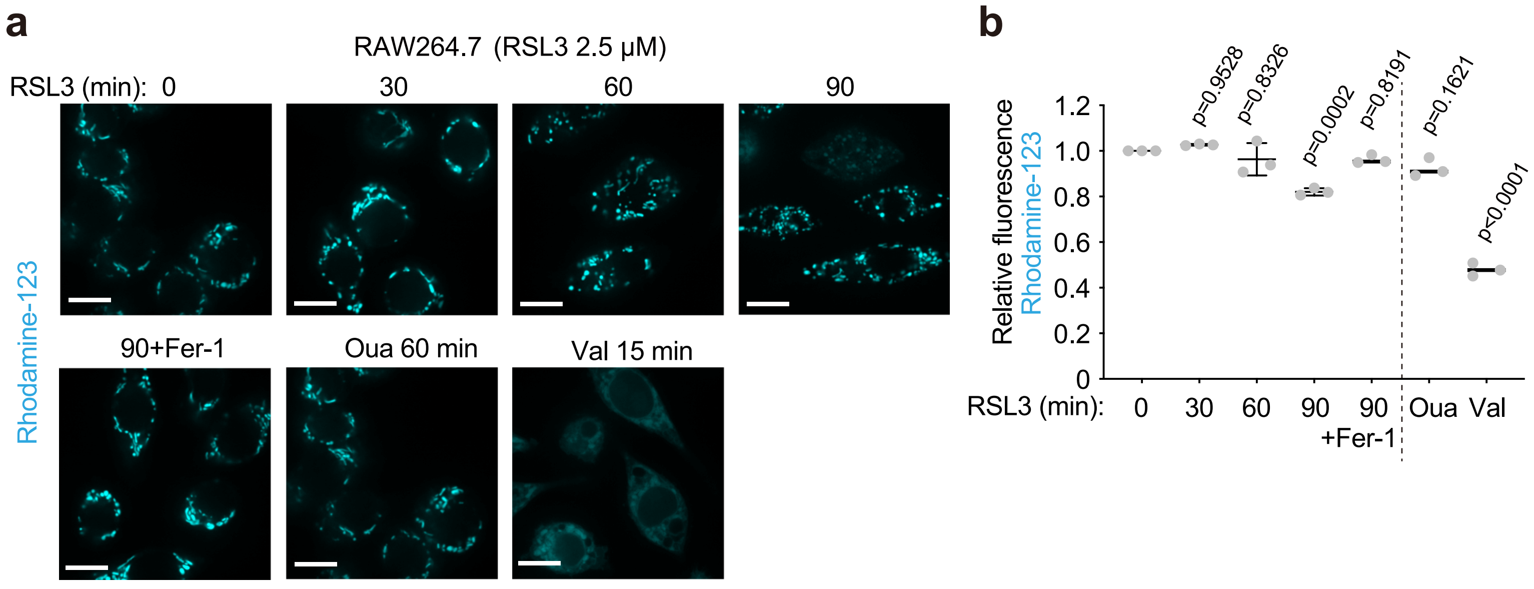

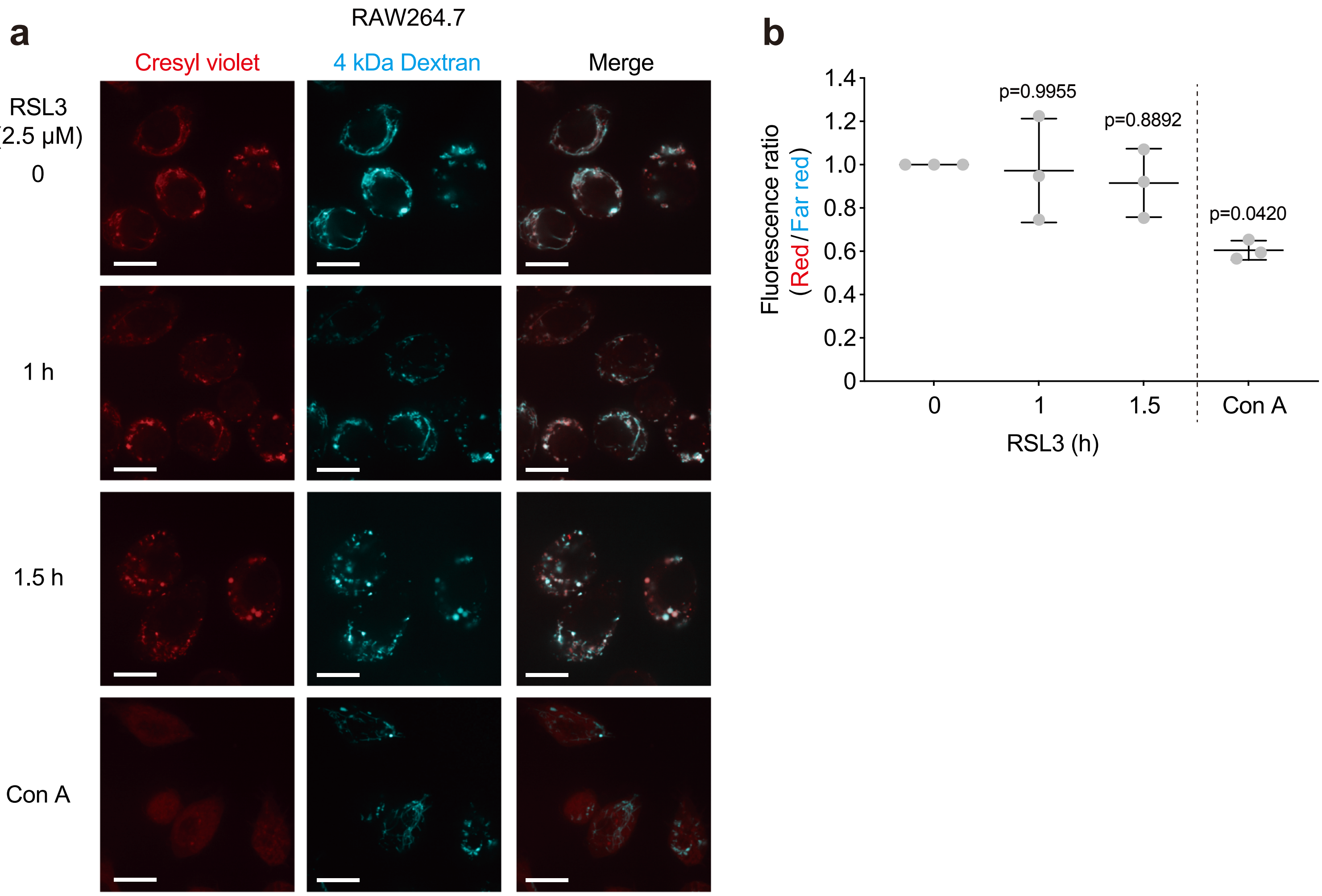

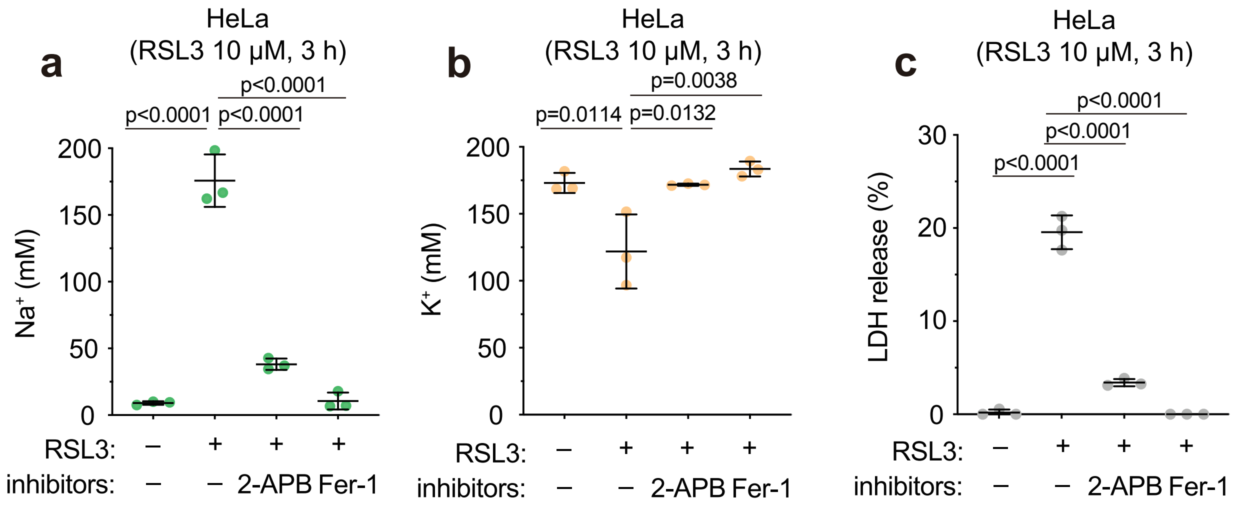
